# The Optimal Combination of Scanning Speed and Fiber-to-Stone Distance for Effective and Efficient Dusting during Holmium: YAG Laser Lithotripsy

**DOI:** 10.1101/2022.06.23.497382

**Authors:** Junqin Chen, Daiwei Li, Wenjun Yu, Zhiteng Ma, Chenhang Li, Gaoming Xiang, Yuan Wu, Junjie Yao, Pei Zhong

**Author notes:** **Corresponding author:** Pei Zhong, PhD, Thomas Lord Department of Mechanical Engineering and Materials Science, Duke University, Box 90300, Durham, NC 27708.

## Abstract

**Objectives:** To investigate mechanistically the effects of fiber scanning speed (*v*_*fiber*_) and fiber tip-to-stone standoff distance (SD) on dusting efficiency during Holmium (Ho): YAG laser lithotripsy (LL)

**Materials and Methods:** Pre-soaked BegoStone samples (23 × 23 × 4 mm, W x L x H) were treated in water using a clinical Ho:YAG laser in dusting mode (0.2 J pulse energy delivered at 20 Hz frequency) at three different SDs (0.10, 0.25 and 0.50 mm) with *v*_*fiber*_ in the range of 0 to 10 mm/s. Stone damage was quantified by optical coherence tomography, bubble dynamics were captured by high-speed imaging, and associated pressure transients were measured using a needle hydrophone. To compare photothermal ablation vs. cavitation in stone dusting, the experiments were further repeated in air (photothermal ablation only), and in water with the fiber tip advanced at a short (0.25 mm) offset distance (OSD) from a ureteroscope to mitigate the bubble collapse toward the stone surface, thus eliminating cavitation-induced damage.

**Results:** Compared to the craters produced by a stationary fiber, the damage troughs produced by a scanning fiber after 100 pulses were significantly larger in volume. The optimal *v*_*fiber*_ for maximum dusting efficiency was found to be 3.5 mm/s for SD = 0.10 ∼ 0.25 mm, resulting in long (17.5 mm), shallow (0.14 – 0.15 mm) and narrow (0.3 – 0.4 mm) troughs. In contrast, the maximum stone damage was produced at an optimal *v*_*fiber*_ of 0.5 mm/s for SD = 0.50 mm, which generates much shorter (2.5 mm), yet deeper (0.35 mm) and wider (1.4 mm) troughs. Greater stone damage was produced in water than in air, especially at *v*_*fiber*_ = 0 – 3.5 mm/s. With the scope end placed near the fiber tip, stone damage could be significantly reduced in water by 29% - 58% for SD = 0.10 mm, by 51% - 82% for SD = 0.25 mm, and by 66% - 100 % for SD = 0.50 mm, compared to those produced without the scope. Together, these findings suggest that cavitation plays an indispensable role in stone dusting by scanning treatment. Moreover, under clinically relevant *v*_*fiber*_ (1 ∼ 3 mm/s), dusting at SD = 0.5 mm (i.e., non-contact mode) may leverage higher frequency of the laser (e.g., 40 to 120 Hz) to harvest the full potential of cavitation damage while significantly reducing the procedure time, compared to its counterpart at SD = 0.1 mm (i.e., contact mode) that promotes photothermal ablation.

**Conclusion:** Dusting efficiency during Ho:YAG LL may be substantially improved by utilizing the optimal combination of *v*_*fiber*_ and SD for a given frequency.

## Introduction

Ureteroscopy with Holmium: YAG (Ho: YAG) laser lithotripsy (LL) has become the first-line therapy for renal calculi over the past decade (1,2). Driven by the development of high-power and high-frequency Ho:YAG lasers, stone dusting has gained clinical popularity over fragmenting because of the shortened procedure time and reduced risk of ureteral damage (3-6). In dusting mode, the Ho:YAG laser is operated at low pulse energy (*E*_*p*_ = 0.2 – 0.5 J) and high pulse repetition frequency (PRF = 12 – 100 Hz). By scanning the laser fiber at various speeds to “paint” over the stone surface line by line and layer by layer, dust-like fragments can be produced that may be discharged spontaneously without the need for basket extraction (5,7,8). The dusting procedure, however, is carried out empirically based on individual urologist’s experience with no consensus on the optimal settings, such as fiber tip-to-stone standoff distance (SD) and fiber scanning speed (*v*_*fiber*_). In addition, the mechanisms of stone dusting produced by a moving fiber during LL remains to be elucidated.

Conventional theory in the past two decades attributes photothermal ablation and microexplosion to be the dominant mechanisms of stone damage in LL (2,9,10). As a result, previous studies often advocate placing the fiber tip in contact with the stone surface, i.e., SD = 0 mm, with the hope to maximize laser energy transmission and treatment efficiency (11-14). Clinically, however, maintaining a close contact of the fiber tip with the stone surface is challenging during LL. In particular, a moving fiber cannot be placed precisely in constant contact with the stone due to either changing surface curvature of the stone (13), the retropulsion effect (15,16), or respiration-induced kidney movement (5). Moreover, a panel of leading endourologists have recently recommended that dusting efficiency can be improved clinically by “defocusing” the laser beam via pulling back the fiber tip slightly away from the stone surface to produce small fragments (5).

Physically, the laser pulse energy deliverable to the stone will decrease significantly even at a short SD (< 1 mm) because of the shallow penetration depth (∼0.4 mm) of the Ho:YAG laser in water (17,18). This unique characteristic of the Ho:YAG laser, coupled with the low *E*_*p*_ used for stone dusting, may diminish the contribution of photothermal ablation (19). Consequently, most of the laser energy in the dusting procedures is likely absorbed by the interposing fluid between the fiber tip and stone surface, leading to the formation of a cavitation bubble with subsequent expansion and collapse (19,20), and temperature rise inside the urinary tract (21,22). Using a short pulse (<100 μs) Ho:YAG laser, we have recently demonstrated that cavitation bubble collapse plays a critical role in stone dusting using a stationary fiber, the efficiency of which can be substantially improved at an optimal SD of 0.5 mm (19).

Several groups have investigated the effect of fiber scanning on stone damage *in vitro* during Ho:YAG LL, mostly in contact mode (i.e., SD ≈ 0 mm where photothermal effects dominate). At *E*_*p*_ = 0.5 J and PRF = 20 Hz, Alhoukhi and colleagues compared two clinically relevant fiber scanning speeds and found that the mass loss was much greater at 3 mm/s than at 1 mm/s (13). Using similar settings, Panthier et al. demonstrated that the maximum stone damage could be produced at *v*_*fiber*_ = 10 mm/s or 2.5 mm/s, compared to lower scanning speeds (23). Both studies used BegoStone phantoms, yet the mechanism that led to maximum efficiency in stone dusting was not investigated. Moreover, two distinctly different damage patterns (i.e., long and continuous trough vs. discrete craters) were produced by either increasing *v*_*fiber*_ from 8.3 mm/s to 25 mm/s (24) or decreasing the number of pulses delivered per scanning distance from 10 pulses/mm to 1 pulse/mm (25). These previous observations suggest that stone damage during scanning treatment may correlate with the overlapping area ratio (OAR) between successive laser pulses, which combines the effect of *v*_*fiber*_ with PRF and fiber diameter.

In this work, we aim to determine the optimal *v*_*fiber*_ at three different SDs (0.10, 0.25, 0.50 mm) under clinically relevant settings for Ho:YAG LL. For a given PRF (i.e., 20 Hz), we vary *v*_*fiber*_ from 0 to 10 mm/s so that a full range of OAR from 0% to 100% can be achieved during scanning, which are correlated with the resultant stone damage. Moreover, we evaluate systematically the contribution of photothermal ablation vs. cavitation to stone dusting by comparison of damage troughs produced in water vs. in air, and with vs. without the use of a ureteroscope that alters the direction of bubble collapse in water. Overall, our results demonstrate that cavitation plays an indispensable role in stone dusting under various SDs, and the treatment efficiency during clinical LL may be significantly improved, with the procedure time greatly shortened, by the optimal combinations of *v*_*fiber*_, PRF and SD.

## Materials and Methods

### Sample preparation protocol

BegoStone samples (BEGO USA, Lincoln, RI) with similar mechanical properties to human kidney stones (26,27) were prepared using 5:2 powder to water ratio by weight (28). Briefly, the mixture was poured into a mold (23 × 23 × 10 mm, LxWxH) and agitated on an orbital shaker (KS 130 control, IKA Works) at 600 RPM for 15 minutes, which localized residual air pockets and impurities in the central region of the sample (28). After a 24-hour curing period, the stone samples were removed from the mold and the two flat surfaces were polished using 1200-grit sandpapers until the sample height was reduced to 4 mm while visible voids were removed from the surface. This preparation procedure allowed us to produce uniform sample surfaces, which improves the consistency of experimental results. All stone samples were soaked in water for 24 hours before LL treatment.

### Scanning fiber experiments

In general, pre-soaked BegoStone phantoms were treated in water at *E*_*p*_ = 0.2 J and PRF = 20 Hz for dusting using a Ho:YAG laser lithotripter (H Solvo 35-watt laser, 270 μm core diameter fiber, NA = 0.26, Dornier MedTech) with a full width at half maximum (FWHM) pulse duration of 70 μs (Fig. 1a). The stones were fixed in a water tank filled with degassed water at room temperature. The laser fiber held by a fiber chuck was positioned perpendicularly to the stone surface, i.e., at 0^°^-laser incident angle, using a 3D positioning stage (VXM-2 step motors with BiSlide-M02 lead screws; Velmex, Bloomfield, NY).

**Figure 1.**
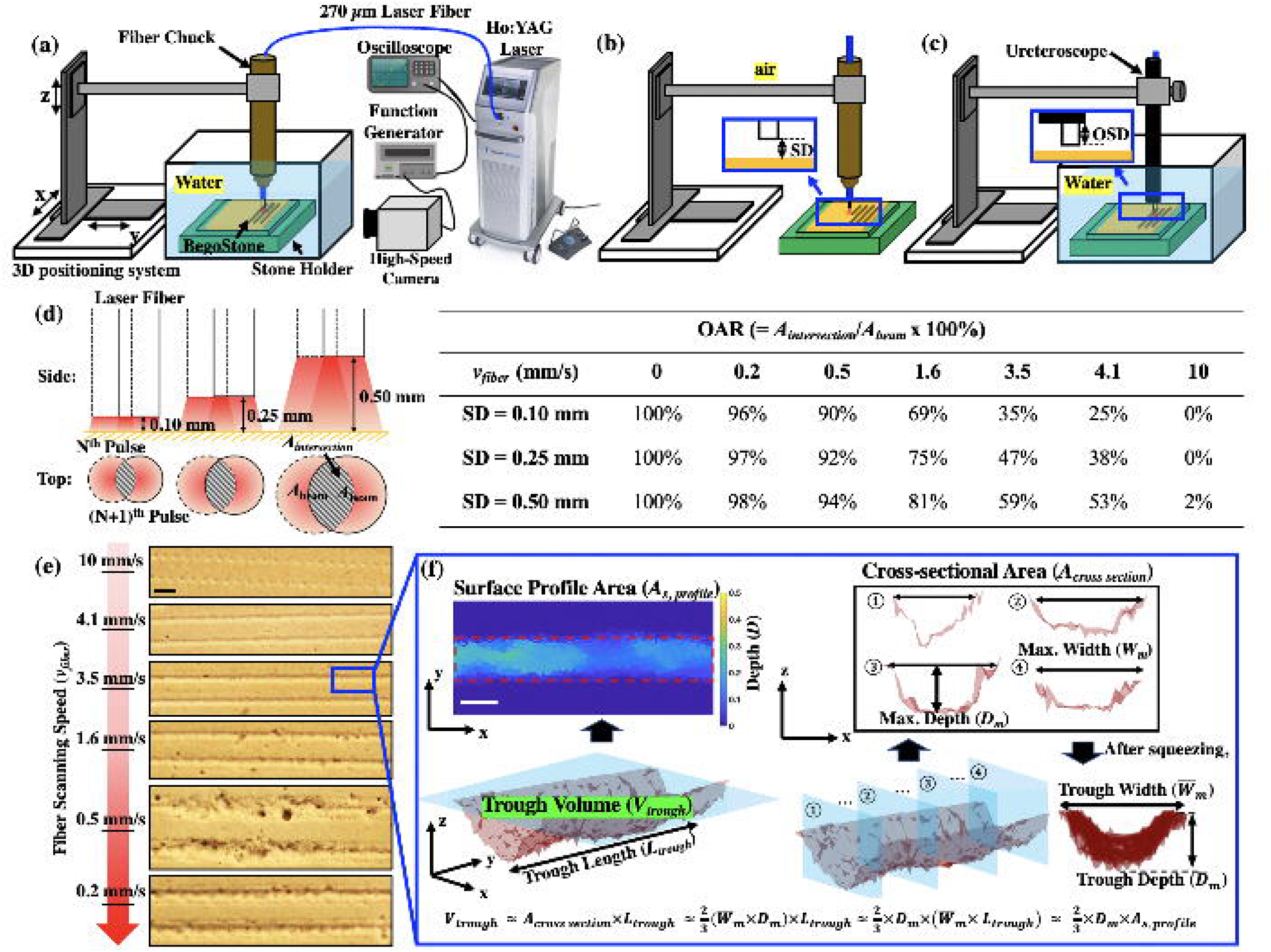
Experimental setup for pre-soaked BegoStone samples treated with perpendicular fiber placed at different fiber tip-to-stone standoff distances (SDs) (**a**) in water synced with high-speed imaging from the side-view, (**b**) in air, and (**c**) in water with a ureteroscope placed at various offset distances (OSDs). A closer view of the gaps between the scope end, fiber tip and stone surface are shown in the blue boxes. (**d**) Overlapping area ratio (OAR) at different SDs and *v*_*fiber*_, which was calculated by Eqn. 2. (**e**) Representative images of stone damage produced at different *v*_*fiber*_ (scale bar = 1 mm). (**f**) An example of a damage trough, which was 3D reconstructed and quantified by OCT scanning including trough volume, surface profile area, trough width and depth (scale bar = 0.5 mm).

A custom MATLAB program (Math-Works, Natick, MA) was used to precisely control the SD (= 0.10, 0.25 and 0.50 mm) between the fiber tip and the sample surface while scanning the fiber across the stone surface at each selected *v*_*fiber*_ (14,25) for creating lines of damage troughs (see Fig. 1e). Following the scan of each line (about 20 mm long), a new fiber tip was prepared using a fiber stripper and cleaved by ceramic scissors. The fiber tip condition was confirmed by inspecting the quality of laser aiming beam on a flat white surface (25,28).

### Quantitative analysis of stone damage

Following LL treatment, the damage troughs on the stone surface were quantified by optical coherence tomography (OCT, OQ Labscope, Lumedica, Durham, NC). The total length of the trough (*L*_*trough*_) produced by 100 pulses was measured, which is proportional to the product of *v*_*fiber*_ and treatment time (5s). Since the maximum dimension quantifiable by our OCT device is 7 × 7 × 2.5 mm (L x W x H), for damage produced at *v*_*fiber*_ > 1 mm/s, we scanned multiple troughs of equal lengths and summed up the results to obtain the total trough volume (*V*_*trough*_). In addition, assuming uniform damage troughs were produced by the Gaussian beam profile of the laser (29) with a mean parabolic cross-sectional area (*Ā*_*cross section*_), *V*_*trough*_ could be estimated by

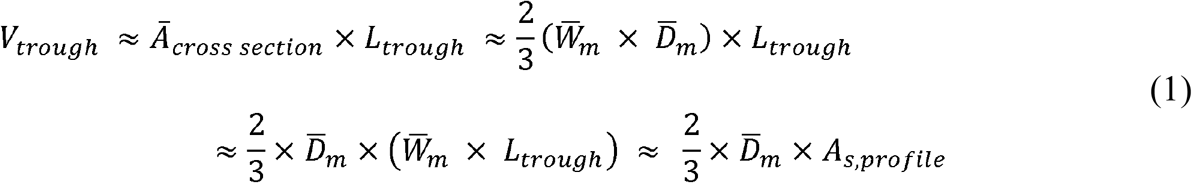

where 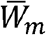 and 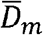 are the means of the maximum width and depth of the cross sections, respectively, and *A*_*s,profile*_ is the surface profile area of the trough. A custom program developed in MATLAB (28) was used to extract these parameters from each damage trough based on the acquired OCT images (see Fig. 1f).

### Assessment of different damage mechanisms

Stone damage in LL could be produced by different mechanisms (2,9,10,19,20,28). To eliminate cavitation-induced stone damage, we performed a second set of experiments in air, which maximizes the laser energy transmission to the stone surface and photothermal ablation (9,28). In addition, we carried out a third set of experiments in water by advancing the fiber tip beyond the distal end of a flexible ureteroscope (Dornier AXIS™, 3.6 F working channel) with a short offset distance (OSD) of 0.25 mm. This method was used to mitigate the bubble collapse toward the stone boundary and thus minimize cavitation-induced damage without affecting the MOSES effect and photothermal ablation of the stone material by the laser pulses in water (19,20,30). Using these strategies, we could compare the contribution of photothermal ablation vs. cavitation damage in stone dusting. For completeness, we also evaluated the effect of the ureteroscope on stone dusting efficiency at two clinically relevant OSDs of 2 mm and 3 mm (31), under the optimal *v*_*fiber*_ at different SDs.

### Effect of the overlapping area ratio (OAR) on dusting efficiency

Stone damage produced by a scanning fiber during LL may depend on the overlap of laser pulses in the irradiated region (24,32). We calculate OAR between two successive laser pulses for a constant *v*_*fiber*_ by (33):

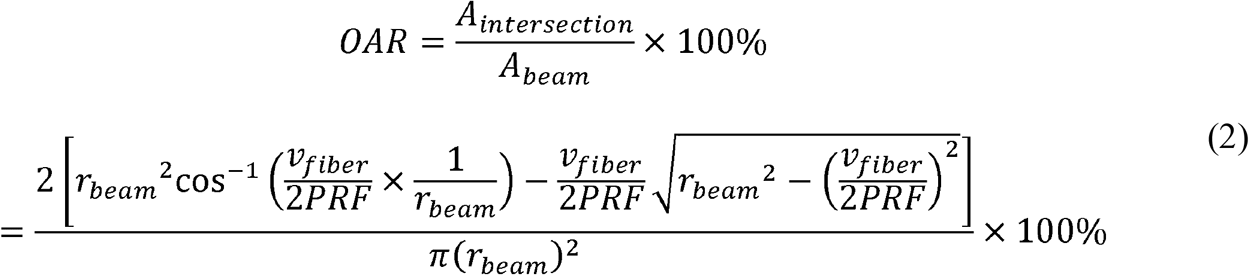

where *A*_*intersection*_ is the intersection area between two laser beams (see Fig. 1d), *A*_*beam*_ is the projected beam area on the stone surface, *r*_*beam*_ is the projected beam radius (0.16 mm, 0.20 mm and 0.27 mm at SD = 0.1, 0.25 and 0.5 mm, respectively) and 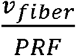 is the inter-pulse distance traveled by the scanning fiber (32). As OAR approaches to 0%, there is no overlap between subsequent pulses, leading to the formation of individual shallow craters on the stone surface (see Fig. 1e at 10 mm/s). In comparison, at OAR of 100%, the laser pulses are fully overlapped with each other, leading to the deep and saturated craters produced by a stationary fiber (19).

### High-speed imaging and pressure transient of bubble dynamics

To capture the bubble dynamics produced in water, we fixed the fiber in position and translated the stone at the selected *v*_*fiber*_ in the opposite direction. A digital time delay generator (BNC 565, Berkeley Nucleonics Corporation) was used to trigger a Kirana5M high-speed video camera (Specialised Imaging, Pitstone, UK) operated at 200,000 frames per second using the internal photodetector signal from the laser (28). Moreover, the bubble-induced pressure transients were measured by placing a needle hydrophone (HNC-1000, Onda, Sunnyvale, CA) at 45^0^ angles, approximately 30 mm from the fiber tip. Furthermore, the experiment was repeated in air to visualize the material ejection from the wet stone surface, using a Phantom v7.3 high-speed video camera (Vision Research, Wayne, NJ) operated at 90,909 frames per second.

### Statistical Analysis

Three-way ANOVA analyses were first carried out to assess the contribution of three factors: treatment condition (in air or in water), *v*_*fiber*_ and SD (0.10 mm, 0.25 mm, and 0.50 mm) on stone dusting efficiency based on the F-test. If any of these F-tests is significant, we then performed a post-hoc test, the Tukey’s Honestly-Significant-Difference (TukeyHSD) test to determine which groups were different from one another. Based on the TukeyHSD test results, we could identify the optimal *v*_*fiber*_ and SD among the various combinations of the test conditions. In addition, we have used two-sample t-tests to compare stone dusting efficiency produced at different OSD levels associated with the flexible ureteroscope.

## Results

### Stone damage produced in water under different scanning speeds and SDs

Figure 2a shows the damage patterns produced under different *v*_*fiber*_ at various SDs in water. Several important features could be observed. First, for a stationary fiber (i.e., *v*_*fiber*_ = 0 mm/s), a small circular crater was formed at SD = 0.10 mm while irregularly shaped craters with enlarged surface profile areas were produced at SD = 0.25 – 0.50 mm. The additional damage around the central crater might be produced by the toroidal bubble collapse following the primary collapse of LL-induced vapor bubble (19). Second, for a scanning fiber, straight damage troughs with varying degrees in 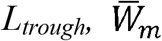 and 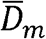 were produced. Notably, when *L*_*trough*_ lengthened significantly at increased *v*_*fiber*_, the corresponding values of 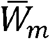 and 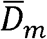 would decrease appreciably in the range of *v*_*fiber*_ = 0.5 mm/s - 10 mm/s (Fig. 2b-2c). Interestingly, 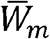 and 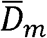 of the craters produced at 0 mm/s were also smaller than their counterparts produced at 0.5 mm/s. Third, at *v*_*fiber*_ = 10 mm/s, individual and shallow circular craters with uniform spacing between them were produced since there was no overlap between two successive fiber positions during scanning. Under such high,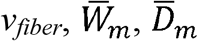, and *A*_*surface*_ (Fig. 2b-d) of the individual craters would all decrease with increasing SD.

**Figure 2.**
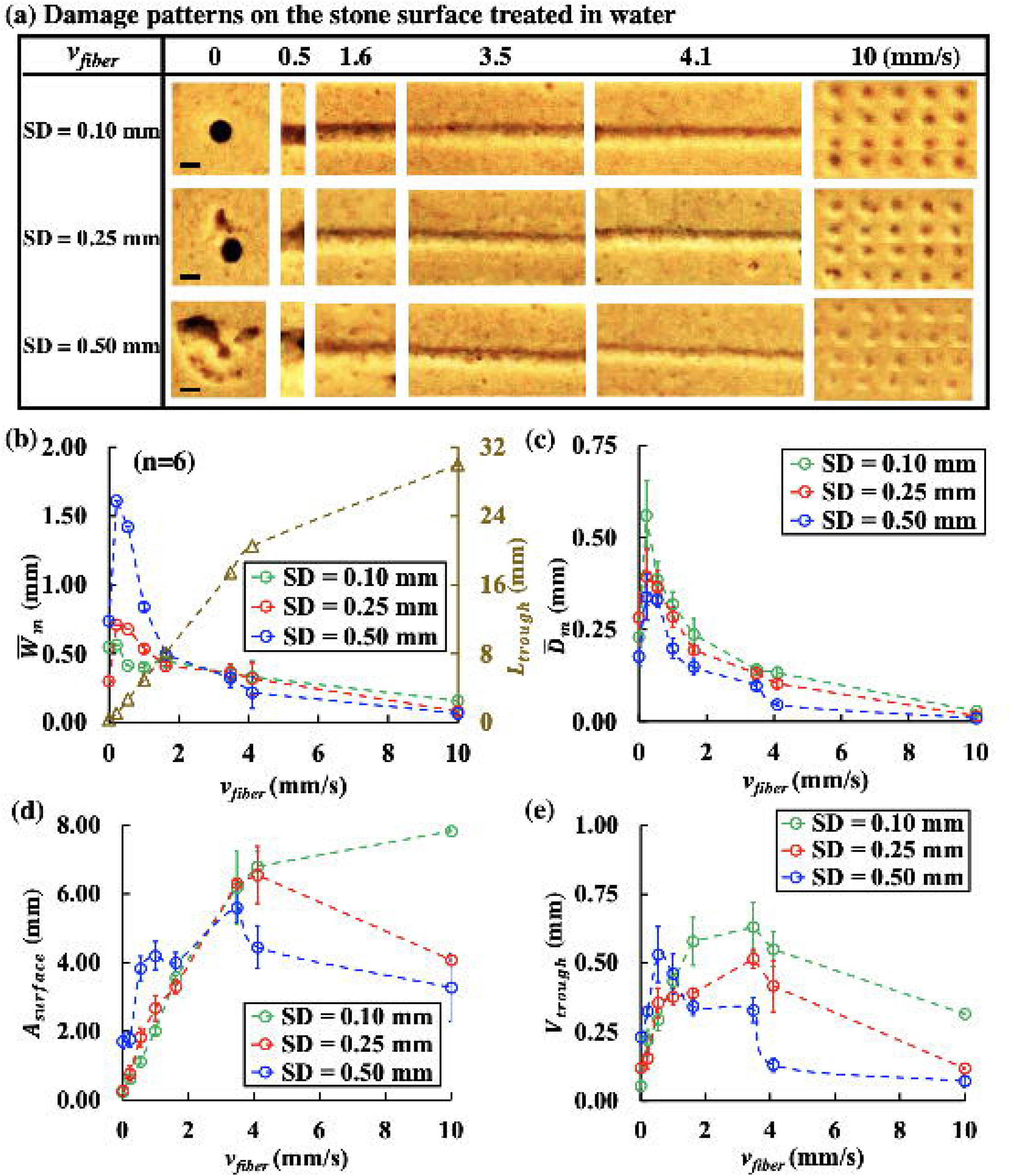
Different damage patterns and characteristic dimensions of the trough produced in water by 100 pulses (0.2 J and 20 Hz) during scanning treatment at various fiber speeds (*v*_*fiber*_). (**a**) Damage craters produced by the stationary fiber (*v*_*fiber*_ = 0 mm/s) and one-fifth of the damage troughs created at different *v*_*fiber*_ under SD = 0.10, 0.25, and 0.50 mm (scale bar = 0.5 mm), (**b**) mean trough width 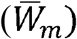 and trough length (*L*_*trough*_), (**c**) mean trough depth 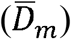, (**d**) surface profile area (*A*_*surface*_), and (**e**) trough volume (*V*_*trough*_) quantified by OCT imaging analysis and plotted vs. *v*_*fiber*_.

More importantly, both *v*_*fiber*_ and SD were found to exert statistically significant influence on *V*_*trough*_ (p<0.001) with the optimal *v*_*fiber*_ for maximum dusting efficiency varying distinctly with SDs (Fig. 2e). At SD = 0.10 mm and SD = 0.25 mm, the trough volumes initially increased with *v*_*fiber*_ and reached their peak values (0.63 mm^3^ and 0.51 mm^3^, respectively) at *v*_*fiber*_ = 3.5 mm/s before decreasing gradually from *v*_*fiber*_ = 4.1 mm/s to 10 mm/s. In comparison, at SD = 0.50 mm, the maximum trough volume (0.53 mm^3^) was produced at an optimal *v*_*fiber*_ = 0.5 mm/s before tapering off thereafter.

### Contribution of photothermal ablation vs. cavitation to stone dusting

To dissect the mechanism of stone dusting under various treatment conditions (*v*_*fiber*_ and SD), we compare stone damage produced in water without or with the flexible ureteroscope at OSD = 0.25 mm, as well as in air.

#### 1) Different bubble dynamics and stone damage characteristics produced by photothermal ablation and cavitation bubble collapse

Figure 3 shows representative high-speed images of the bubble dynamics produced by the laser-fluid-stone interaction at different SDs using a stationary fiber. Without the scope, the maximum bubble size formed was found to increase with SD from 0.10 mm to 0.50 mm as more laser pulse energy was absorbed by the interposing fluid between the fiber tip and stone surface (19). Consequently, the strongest pressure transient was generated by the bubble collapse at SD = 0.50 mm (Fig. 3a). With the scope at OSD = 0.25 mm, although the bubble expansion in relation to laser transmission to the stone (i.e., the MOSES effect) were not affected, the collapse of the bubble was distracted by the proximity of the scope tip and moved away from the stone surface (Fig. 3b). When the bubble collapse was mitigated by the scope tip, the resultant craters became much smaller and shallower, compared to their counterparts produced without the scope at different SDs (Fig. 3c). Furthermore, stone damage produced in air (without cavitation) was similar to the small circular craters produced in water with the scope, except that the sizes of the central crater and surrounding burn mark would decrease with increasing SD (Fig. 3c).

**Figure 3.**
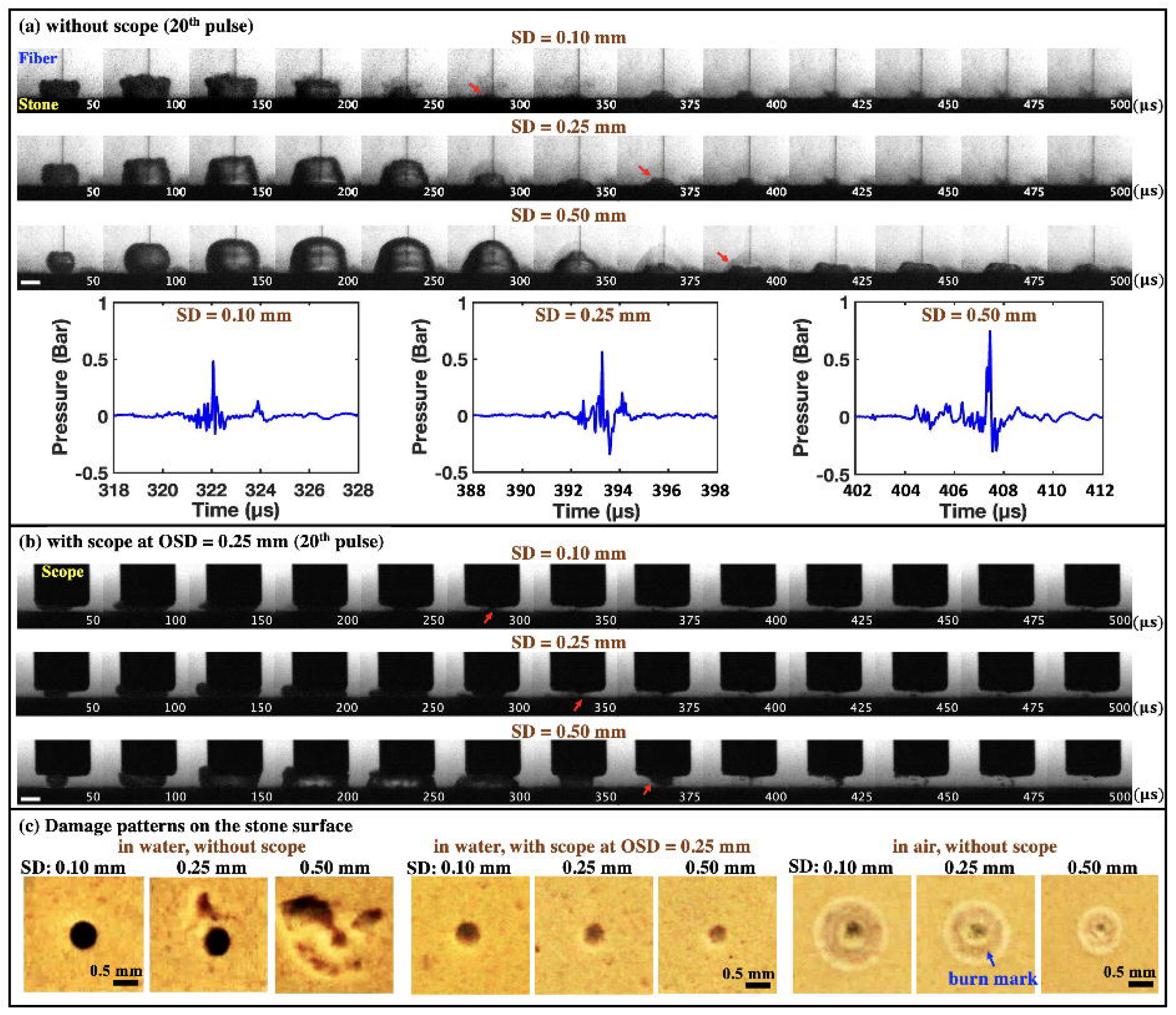
Representative high-speed imaging sequences of bubble dynamics produced in water near the BegoStone surfaces during dusting (*E*_*p*_ = 0.2 J and PRF = 20 Hz) using a stationary fiber. (**a**) without the ureteroscope (or scope) at SD = 0.10 mm, 0.25 mm, and 0.50 mm and the corresponding pressure transients measured by the needle hydrophone, and (**b**) with the scope at OSD = 0.25 mm (scale bar = 1 mm). In both (**a**) and (**b**), the red arrows indicate the direction of bubble collapse under different treatment conditions. (**c**) Damage patterns produced on the BegoStone surfaces after 100 pulses in water without and with the scope, and in air (scale bar = 0.5 mm). The blue arrow indicates the burn mark around the central craters.

#### 2) Cavitation bubble collapse toward the stone surface is indispensable for effective stone dusting

In general, *V*_*trough*_ produced in water without the scope is significantly larger than its counterpart with the scope (Fig. 4), confirming the indispensable role of cavitation bubble collapse in stone dusting (19) even during scanning treatment. Although the variations of damage patterns with *v*_*fiber*_ at different SDs were similar, the largest reduction in stone damage by the proximity of the scope tip was 5.3-fold, observed at SD = 0.50 mm under the optimal *v*_*fiber*_ of 0.5 mm/s. In comparison, the maximum reduction in stone damage at SD = 0.10 mm and SD = 0.25 mm were 1.5- and 2.5-fold, respectively, observed under a different optimal *v*_*fiber*_ of 3.5 mm/s. These significant reductions in *V*_*trough*_ may be attributed to the substantial diminishments in both 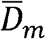 and 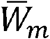 because of the suppressed bubble collapse, especially for SD = 0.50 mm (Fig. 4e to 4l).

**Figure 4.**
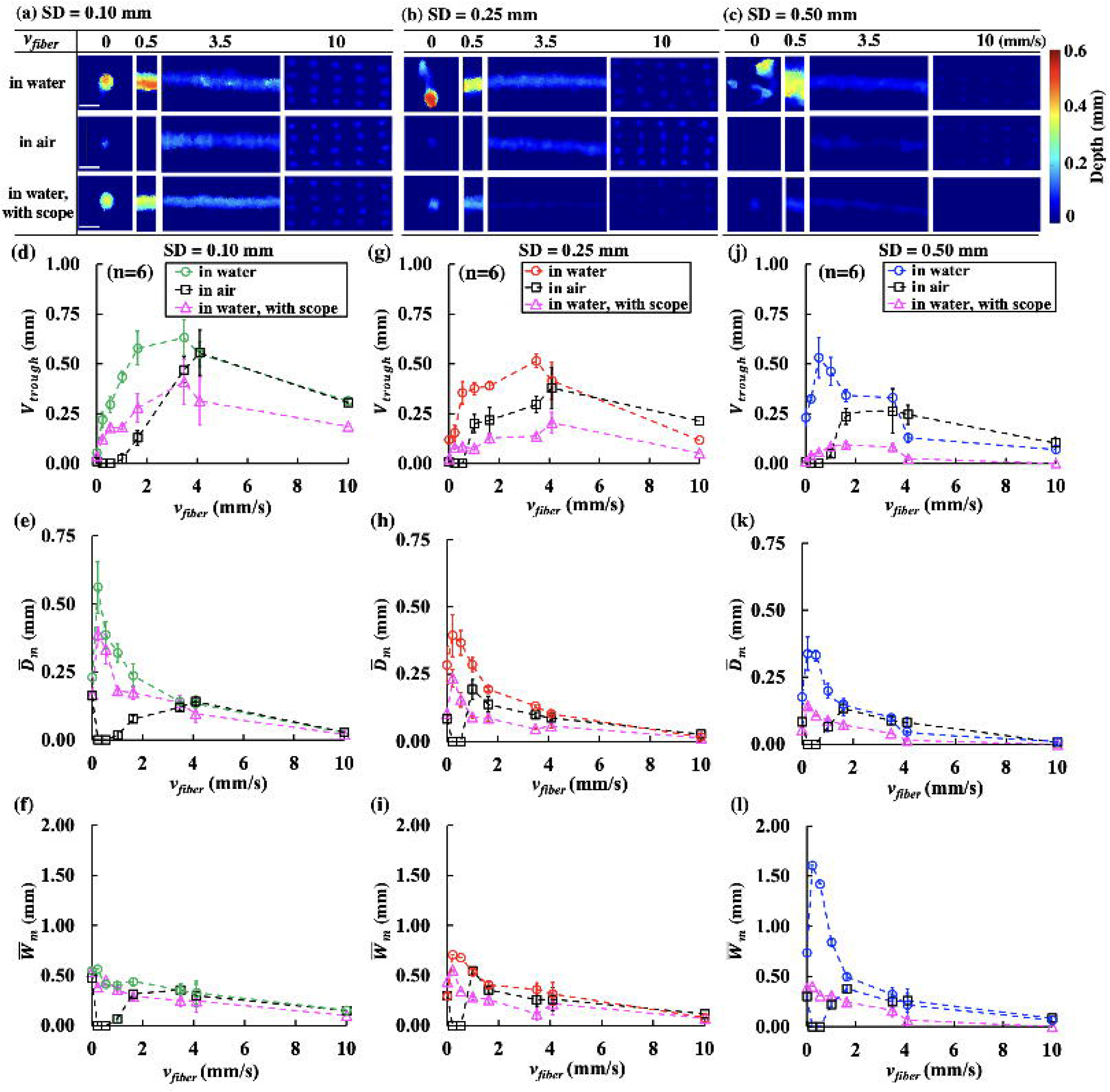
OCT reconstruction of damage patterns on BegoStone surfaces produced by 100 pulses (0.2 J and 20 Hz) in water and in air without scope, and in water with scope at OSD = 0.25 mm delivered at (**a**) SD = 0.10 mm, (**b**) SD = 0.25 mm, and (**c**) SD = 0.50 mm (scale bar = 1 mm). Trough volume *(V*_*trough*_) (**d, g, j**), mean trough depth 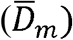 (**e, h, k**) and mean trough width 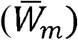 (**f, i, l**) are plotted vs. *v*_*fiber*._

Even when cavitation was eliminated with more laser energy delivered to the stone in air (9,10,28), the resultant stone damage was still found to be less than its counterpart produced in water without the scope for *v*_*fiber*_ < 4.1 mm/s, regardless of SD. These findings confirm again that besides photothermal ablation, cavitation contributes vitally to stone dusting in Ho:YAG LL (19).

#### 3) Effects of scanning speed and SD on bubble dynamics and resultant acoustic emission

Representative high-speed imaging frames of bubble expansion and collapse at different SDs and *v*_*stone*_ are shown in Figure 5a. To facilitate high-speed imaging, the camera and the fiber were fixed during the experiment while the stone was translated at different speeds (*v*_*stone*_, from right to left) to mimic the scanning treatment. The shape of the bubble expansion was significantly altered by the movement of the stone, especially at short SD (e.g., 0.10 mm) and slow *v*_*stone*_ (e.g., 0.5 mm/s). At low *v*_*stone*_ (=0.5 mm/s), the maximum bubble size increased appreciably compared to its counterpart at high *v*_*stone*_ (=10 mm/s) regardless of SDs. This difference is likely caused by the more rapid growth of the damage trough produced at a slow scanning speed, leading to greater 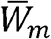, and 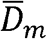 (Fig. 4e to 4l) and thus increased laser absorption in the interposing fluid than those produced at a fast scanning speed. Concomitantly, the highest acoustic pressure emitted by the bubble collapse was measured at SD = 0.50 mm, followed by SD = 0.25 mm, with the lowest acoustic emission produced at SD = 0.10 mm (Fig. 5a). Furthermore, the peak pressure varied significantly with SD and statistical differences were observed in the acoustic emission between *v*_*stone*_ ≤ 4.1 mm/s and *v*_*stone*_ = 10 mm/s (p < 0.04). These results are consistent with the general notion that the effect of cavitation in stone dusting will increase with SD, reaching a maximum at the optimal SD of 0.50 mm (Fig. 4).

**Figure 5.**
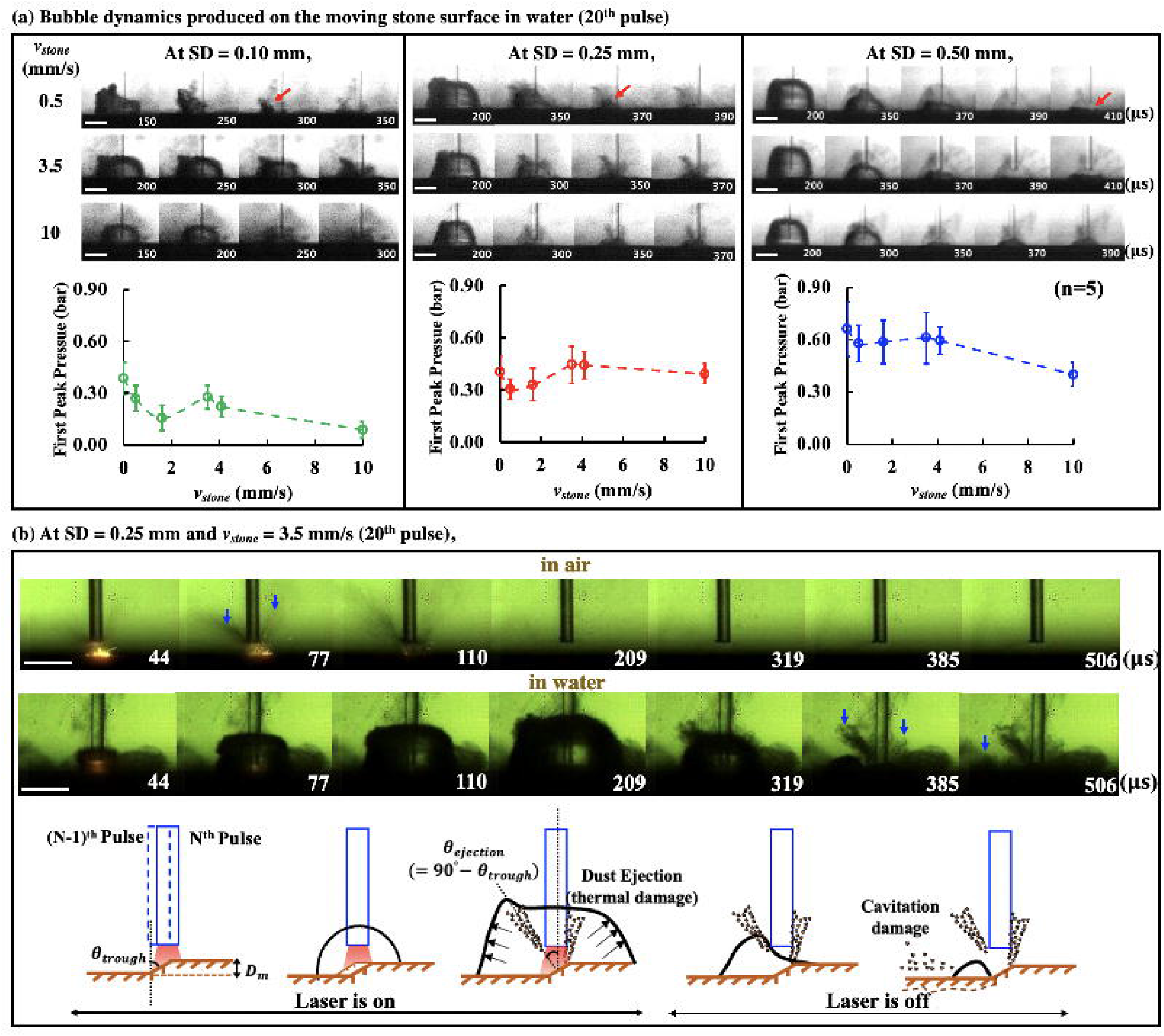
(**a**) High-speed imaging sequences in which the bubble produced on the stone surface moving at different speeds (*v*_*stone*_) expands to its maximum volume and collapses at different SDs (scale bar = 1 mm). The red arrows indicate the direction of bubble collapse. The first peak pressure obtained from the hydrophone measurements is plotted vs. *v*_*stone*_. The direction of stone movement in the high-speed imaging was from right to left. (**b**) Representative high-speed imaging sequences of laser-stone interaction in air vs. in water at one of the optimal settings (SD = 0.25 mm, *v*_*fiber*_ = 3.5 mm/s) (scale bar = 1 mm) and the sketch of general process of stone dusting produced in water, where *θ*_*trough*_ is the angle of trough damage produced by the first N pulse and *θ*_*ejection*_ is the angle of material ejection from stone surface. The blue arrows indicate the material ejection.

Figure 5b shows an example of the detailed physical processes involved in air vs. in water during scanning treatment under one of the optimal settings (SD = 0.25 mm, *v*_*fiber*_ = 3.5 mm/s). The laser-stone (in air) or laser-fluid-stone (in water) interaction during scanning treatment created by the N^th^ pulse was influenced by the surface damage already produced by the previous (N-1) pulses. As such, while portion of the laser pulse energy irradiated (as the fiber moved equivalently from left to right) was absorbed by the trailing damage trough surface, the rest of the energy would be absorbed by the untreated surface in front of the scanning fiber (toward its right side). As a result, two distinctly different directions of material ejection could be observed in front and behind the scanning fiber (see arrows at 77 μs in Fig. 5b). In general, the central direction of material ejection caused by photothermal ablation is approximately perpendicular to the stone surface (15,34), which is consistent with our observation in air.

In comparison, the vapor bubble created in water during scanning was significantly distorted (especially at low speed and short SD, see Fig. 5a) compared to the bubble produced by a stationary fiber (Fig. 4a). The bubble dimension was larger behind (left) than in front of the fiber (right), presumably due to the stronger laser absorption in the interposing fluid in the damage trough behind the fiber, supplemented by the material ejection and laser-dust interaction (i.e., flashes inside the expanding bubble at 44 μs in Fig. 5b). Following the cessation of the laser pulse and the maximum expansion, the bubble tended to collapse toward the left into the damage trough (see the sketch in Fig. 5b). This asymmetric expansion and collapse of the bubble produced by a scanning fiber (see Video 1) may cause additional material removal by the multi-foci collapse of the bubble (see arrows at 385 μs and 506 μs in Fig. 5b) following the initial photothermal damage.

#### 4) Effect of fiber tip OSD from the ureteroscope on stone dusting efficiency

When the bubble collapse to the stone surface was mitigated by the scope tip at OSD =M0.25 mm (Fig. 4d-4j), the resulting *V*_*trough*_ was reduced by 29% - 58% for SD = 0.10 mm, 51% - 82% for SD = 0.25 mm, and 66% - 100 % for SD = 0.50 mm in the range of *v*_*fiber*_ = 0 – 10 mm/s (Fig. 6a). These results clearly demonstrate the critical role that cavitation bubble collapse plays in stone dusting, especially for SD = 0.25 and 0.50 mm at a slow scanning speed (i.e., *v*_*fiber*_ ≤ 3.5 mm/s). Clinically, the OSD is typically in the range of 2-3 mm (31). Under such conditions, the presence of the ureteroscope tip on stone damage was found to be statistically insignificant under the optimal speed *v*_*fiber*_ = 3.5 mm/s for SD = 0.10 mm or 0.25 mm (p > 0.4) (Fig. 6b). Only under SD = 0.50 mm with its optimal speed *v*_*fiber*_ = 0.5 mm/s, the reduction of *V*_*trough*_ at OSD = 2 or 3 mm was 67% or 38%, which was statistically significant (p<0.01).

**Figure 6.**
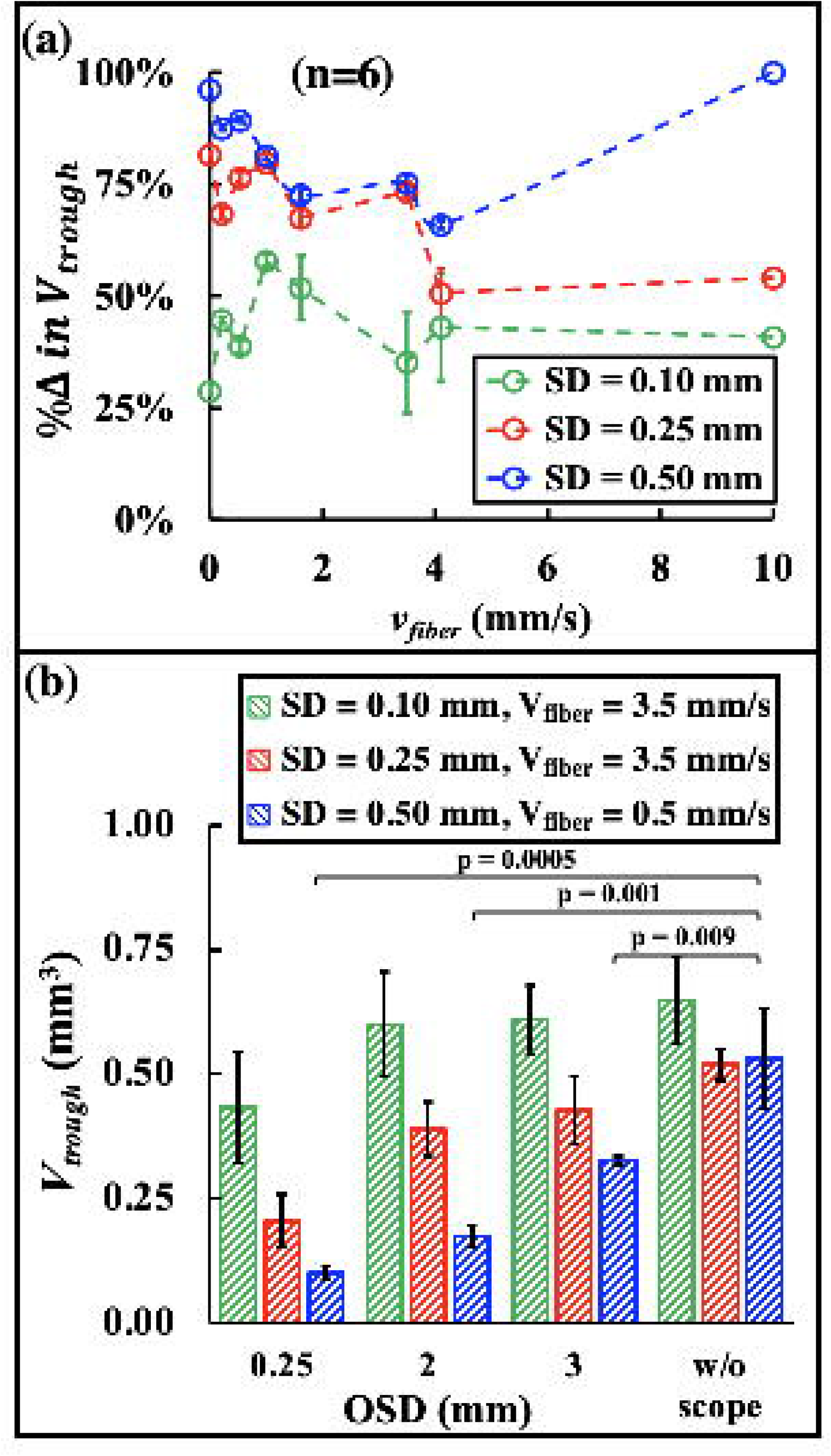
(**a**) The percentage of reduction in trough volume (%Δ in *V*_*through*_) at different SDs and *v*_*fiber*_ were calculated by %Δ where 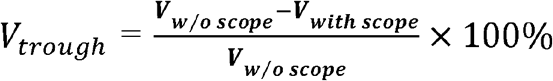 is the *V*_*w/oscope*_ trough volume produced by the treatment without the scope and *v*_*with scope*_ is the trough volume produced by the treatment with the scope at OSD = 0.25 mm. (**b**) Trough volume vs. OSDs under the three optimal conditions: SD = 0.10 mm at *v*_*fiber*_ = 3.5 mm/s, SD = 0.25 mm at *v*_*fiber*_ = 3.5 mm/s, and SD = 0.50 mm at *v*_*fiber*_ = 0.5 mm/s.

## Discussion

The introduction of high-power (up to 120 W) and high-frequency (up to 120 Hz) Ho:YAG lasers in recent years have fundamentally changed the mode of LL from fragmenting to dusting, pop-dusting, or popcorning (5,8). These new treatment modes significantly reduce the overall procedure time, eliminate the need for stent use while minimizing ureteral damage in LL (35,36). Despite these advances, fundamental challenges still exist in LL to improve stone dusting efficiency via optimization of treatment strategy (37-40). In this work, we have demonstrated that the efficiency of stone dusting can be maximized by an optimal combination of *v*_*fiber*_ and SD produced by a short pulse Ho:YAG laser at 20 Hz PRF.

The overarching goal in stone dusting is to create fine fragments (e.g., < 1 mm) while maximizing the material removal (8,16). Dusting efficiency may be influenced by multiple settings and procedural parameters in LL, including *E*_*p*_, PRF, SD, and *v*_*fiber*_, in addition to stone composition, shape and size (16,41-43). Clinically, it has been advocated that dusting should be performed using the lowest possible *E*_*p*_ to create fine fragments while minimizing the undesirable retropulsion effect (16). Moreover, careful adjustment of the fiber tip to stone surface distance (or SD) has been recommended for clinical LL (5), and its influence on treatment outcome demonstrated recently in laboratory studies (19,28). Furthermore, scanning the fiber tip over the stone surface may avoid saturation in stone damage when laser pulses are delivered to a fixed spot (23,44). However, the optimal *v*_*fiber*_ and SD have not been determined, nor has the associated mechanism of action been elucidated. In this study, using high-speed imaging, hydrophone measurement, and the proximity effect of the ureteroscope tip, we shed new light on the physical processes and critical parameters in LL that govern the treatment outcome.

Our results suggest that maximum material removal during dusting can be achieved by an optimal combination of the three characteristic dimensions in the damage trough (i.e., 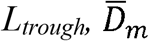 and 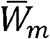). Importantly, the optimal treatment conditions and associated mechanism of action depend critically on SD and the OAR between successive laser pulses (Fig. 7). In contact mode (i.e., SD = 0.10 mm) when photothermal ablation dominates, the maximum dusting efficiency is produced at an optimal *v*_*fiber*_ of 3.5 mm/s, corresponding to an OAR of 35%. Under such treatment conditions, *V*_*trough*_ will increase rapidly with *v*_*fiber*_ and reach a maximum at *v*_*fiber*_ = 3.5 mm/s before tapering off thereafter. This optimization process is driven primarily by the significant increase of *L*_*trough*_ (see Fig. 4a) before the concomitantly reduced 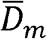 and 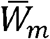 diminish markedly (see Fig. 4e and 4f).

**Figure 7.**
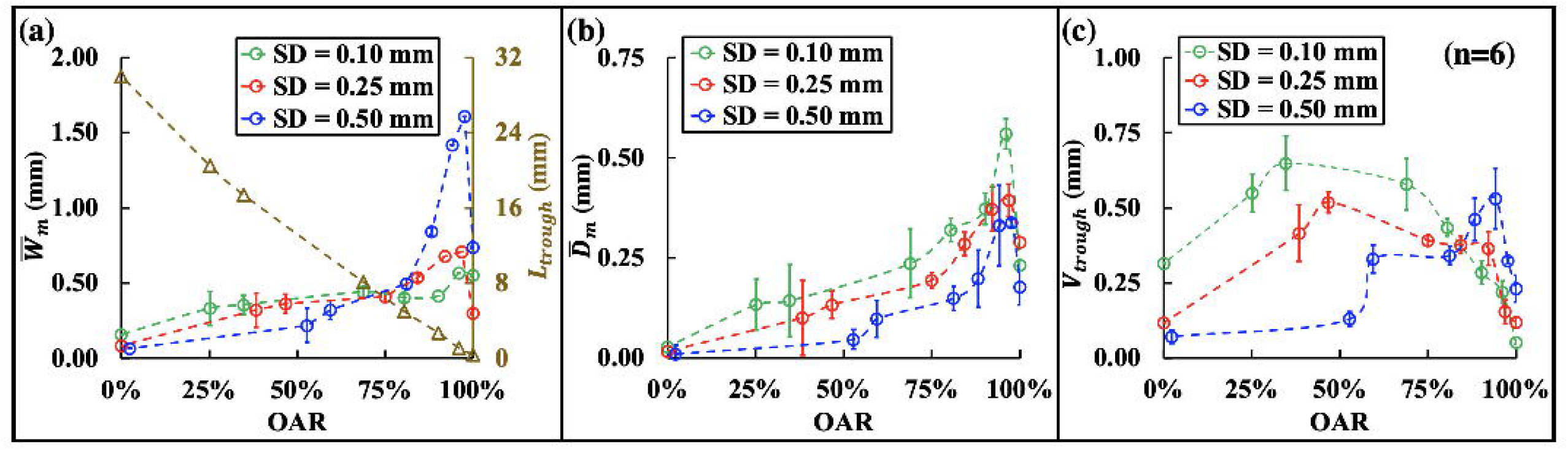
(**a**) Length (*L*_*trough*_) and mean width 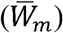, (**b**) mean depth 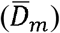, and (**c**) volume (*V*_*trough*_) of the damage trough vs. the overlapping area ratio (OAR) between the successive laser pulses.

In contrast, in non-contact mode (e.g., SD = 0.50 mm) when cavitation-enhanced stone dusting is maximized (19) while photothermal effect is substantially reduced, the maximum dusting efficiency is produced at a much slower optimal *v*_*fiber*_ of 0.5 mm/s, corresponding to a significantly increased OAR of 94%. Under such treatment conditions, the significant overlap between successive laser pulses will greatly enhance the accumulative effect of cavitation damage with substantially broadened 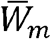 and deepened 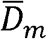 (see Fig. 4k and 4l) despite that the increase in *L*_*trough*_ is limited (see Fig. 4c).

Although the significant contribution of cavitation to stone dusting has been recently demonstrated using fixed spot treatment with up to 640 pulses (19), there is concern regarding whether such a critical observation will hold true under clinically relevant LL conditions (39,40). As such, it is important to note that when the contribution of bubble collapse was mitigated (Fig. 6a), stone dusting efficiency could be greatly reduced (51% to 100%) during scanning at various *v*_*fiber*_ from SD = 0.25 mm to 0.5 mm. Even at SD = 0.1 mm when a strong photothermal effect was anticipated, a remarkable decrease in stone damage (29% to 58%) could still be observed by altering bubble collapse.

Altogether, these results imply an indispensable role of cavitation in stone dusting from every single bubble collapse. As captured by the high-speed imaging in Fig. 5, the bubble collapse may help removal of dust or damaged substances from the trough surface, and thus enhance the energy transmission of subsequent laser pulses to reach the underneath stone material for effective photothermal ablation. Such a scenario may present an alternative mechanism by which the collapse of cavitation bubbles, even under suboptimal conditions (i.e., contact mode or SD = 0.1 mm), may contribute to stone damage. In contrast, using non-contact mode at the SD conductive for cavitation damage (e.g., SD = 0.5 mm), the dynamics of bubble collapse will be critically influenced by the topology of the stone surface and proximity to the scope tip (Fig. 5 and Fig. 6b). Further studies are warranted to elucidate the mechanism of action for cavitation-driven stone dusting under clinically relevant treatment conditions and inside the kidney tissue environment (40).

Moreover, it is worth noting when *v*_*fiber*_ increases from 0.2 mm/s to 10 mm/s, the significant downward trends in 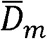 and 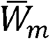 observed in Fig. 4 are much more pronounced in water without the scope (i.e., strong bubble collapse) than in water with the scope at OSD = 0.25 mm (i.e., mitigated bubble collapse). Therefore, without bubble collapse a higher *v*_*fiber*_ is needed to produce a large *L*_*trough*_ that compensates the reduction in 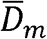 and 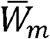 in order to maximize *V*_*trough*_. This feature can be seen from the distinct shift of the optimal *v*_*fiber*_ from 3.5 mm/s to 4.1 mm/s under SD = 0.25 mm, and from 0.5 mm/s to 3.5 mm/s under SD = 0.50 mm when the bubble collapse toward the stone surface was eliminated. This observation is consistent with previous studies, in which a faster scanning speed (e.g., 3 mm/s vs. 1 mm/s) was found to produce higher dusting efficiency regardless of PRF in contact mode (13). Under similar settings (0.5 J, 20 Hz) using another laser lithotripter with different pulse profiles, the maximum dusting efficiency could be achieved at even higher *v*_*fiber*_ of 10 mm/s (23). However, it should be noted that slow *v*_*fiber*_ is much easier to manipulate during clinical LL and preferable over fast scanning speeds to avoid unintentional ureteral wall injury (23).

The clinical significance of our results can be further explored by examining the optimal PRF for a given *v*_*fiber*_. Physically, the characteristics in the stone damage trough are determined by the OAR (Fig. 7), which unifies *v*_*fiber*_ and PRF into a ratio correlating with the inter-pulse distance (see Eq. 2). Therefore, the same OAR between two successive laser pulses during scanning will ensure similarities in laser-fluid-bubble-stone interaction and resultant dusting damage, independent of the specific values of *v*_*fiber*_ and PRF. Consequently, we can estimate the optimal PRF under clinically relevant *v*_*fiber*_ within the range of 1 mm/s to 3 mm/s (13). The results, summarized in Table 1 together with the procedure time, suggest that the optimal PRF for stone dusting in contact mode (SD = 0.10 mm) will vary from 6 Hz to 17 Hz, which is within the frequency range for conventional low-power Ho:YAG lasers (45). In comparison, the optimal PRF for stone dusting using non-contact mode (SD = 0.50 mm) will be much higher, increasing from 40 Hz to 120 Hz, which can only be produced by contemporary high-power/high frequency Ho:YAG lasers. If operating the contact mode in such a high frequency range, the dusting efficiency will decrease by 56% - 66% at 1 mm/s, 33% - 61% at 2 mm/s or 11% - 56% at 3 mm/s

**Table 1.** Comparison of the optimal pulse repetition frequency (PRF) and procedure time under three clinically relevant fiber speeds (*v*_*fiber*_): 1 mm/s, 2 mm/s and 3 mm/s at a fiber-to-stone standoff distance (SD) of 0.10 mm (contact mode) and 0.50 mm (non-contact mode).

| **SD = 0.10 mm (contact mode)** |  |  |  |
| --- | --- | --- | --- |
| Optimal $v_{\text{fiber}}$ at PRF = 20 Hz | Optimal PRF at different $v_{\text{fiber}}$ | | |
|  | 1 mm/s | 2 mm/s | 3 mm/s |
| 3.5 mm/s ($\frac{v_{\text{fiber}}}{\text{PRF}} = 0.175$) | 6 Hz | 11 Hz | 17 Hz |
| Time required for 100 pulses, $t_1$ | 17.5 s | 8.7 s | 5.8 s |
| **SD = 0.50 mm (non-contact mode)** |  |  |  |
| 0.5 mm/s ($\frac{v_{\text{fiber}}}{\text{PRF}} = 0.025$) | 40 Hz | 80 Hz | 120 Hz |
| Time required for 100 pulses, $t_2$ | 2.5 s | 1.3 s | 0.8 s |
| Ratio of Treatment time, $t_1 : t_2$ | 7 : 1 | | |

Furthermore, it is important to note that the procedure time decreases with increased PRF. Using the respective optimal PRFs, the procedure time for the non-contact mode will be significantly shorter (1/7) compared to its counterpart for the contact mode under clinically viable scanning speeds (1 to 3 mm/s). All in all, stone dusting via non-contact mode at an optimal SD (e.g., 0.50 mm) will maximize cavitation damage that offers competitive treatment efficiency per pulse achievable at much higher PRF and thus greatly reduced procedure time, compared to the conventional photothermal ablation treatment strategy via contact mode (e.g., SD = 0.10 mm). It should be noted that BegoStone phantoms are used in this study. Future work using human renal calculi and in kidney-like tissue environment are warranted to determine the optimal stone dusting strategy *in vivo*.

## Conclusion

In this study, we have investigated the effects of fiber scanning speed and SDs on stone dusting during Ho:YAG LL. Our results suggest that dusting efficiency during scanning treatment can be significantly improved by the optimal combinations of *v*_*fibe*r_, PRF and SD. Moreover, the substantial reduction in stone damage by mitigating bubble collapse toward the stone surface via the scope proximity clearly demonstrates the indispensable role of cavitation in stone dusting at various SDs. Most importantly, compared to the conventional photothermal ablation treatment strategy via the contact mode (e.g., SD = 0.10 mm), the non-contact mode at an optimal SD = 0.50 mm may offer a competitive treatment option for effective and efficient stone dusting leveraged by maximizing cavitation damage under high PRF with shortened procedure time.

## Acknowledgements

The authors are grateful to Dornier *MedTech* for providing the H Solvo laser used in this study. They would also like to thank Dr. Michael Lipkin, Dr. Glenn Preminger, and Dr. Zachary Dionise for their gracious accommodation for engineering students to observe clinical laser lithotripsy procedures at Duke University Medical Center, their education about kidney stone disease, and valuable feedback throughout this research project. This study was one of the three research projects in a new graduate course entitled “Engineering Technology in Urology Applications”, designed specifically to foster the research interest and career development of engineering students in benign urology, and to promote interdisciplinary collaboration between engineering and urology. This work is supported by the National Institute of Health (NIH) through grants 5P20 DK123970-02 and 2R01DK052985-24A1.

